# Kv4.1, a key ion channel for low frequency firing of dentate granule cells, is crucial for pattern separation

**DOI:** 10.1101/670018

**Authors:** Kyung-Ran Kim, Seung Yeon Lee, Yoonsub Kim, Min Jeong Kwon, Hyeon-Ju Jeong, Sanghyeon Lee, Young Ho Suh, Jong-Sun Kang, Hana Cho, Suk-Ho Lee, Myoung-Hwan Kim, Won-Kyung Ho

**Affiliations:** Department of Physiology, Seoul National University College of Medicine, 103 Daehak-ro, Jongno-gu, Seoul 03080, Korea; Neuroscience Research Institute, Seoul National University College of Medicine, 103 Daehak-ro, Jongno-gu, Seoul 03080, Korea; Department of Biomedical Science, Seoul National University College of Medicine, 103 Daehak-ro, Jongno-gu, Seoul 03080, Korea; Department of Molecular Cell Biology, Sungkyunkwan University School of Medicine, Suwon, Korea; Department of Physiology, Sungkyunkwan University School of Medicine, Suwon, Korea

**Keywords:** Kv4.1, dentate gyrus, mature granule cell, low excitability, pattern separation

## Abstract

The dentate gyrus (DG) in the hippocampus may play key roles in remembering distinct episodes through pattern separation, which may be subserved by the sparse firing properties of granule cells (GCs) in the DG. Low intrinsic excitability is characteristic of mature GCs, but ion channel mechanisms are not fully understood. Here, we investigated ionic channel mechanisms for firing frequency regulation in GCs of the rat hippocampus, and identified Kv4.1 as a key player. Immunofluorescence analysis showed that Kv4.1 was preferentially expressed in the DG, and its expression level was higher at 8-week than 3-week-old mice, suggesting a developmental regulation of Kv4.1 expression. With respect to firing frequency, GCs are categorized into two distinctive groups: low frequency (LF) and high frequency (HF) firing GCs. Input resistance (Rin) of most LF-GCs is lower than 200 MΩ, suggesting that LF-GCs are fully mature GCs. Kv4.1 channel inhibition by intracellular perfusion of Kv4.1 antibody increased firing rates and gain of the input-output relationship selectively in LF-GCs with no significant effect on resting membrane potential and Rin, but had no effect in HF-GCs. Importantly, mature GCs from mice depleted of Kv4.1 transcripts in the DG showed increased firing frequency, and these mice showed an impairment in contextual discrimination task. Our findings suggest that Kv4.1 expression occurring at late stage of GC maturation is essential for low excitability of DG networks and thereby contributes to pattern separation.

## Introduction

Learning similar events rapidly as distinct episodic memories and retrieving them as such, a computational process called *pattern separation*, are the hallmarks of hippocampal functions. It is postulated that the dentate gyrus (DG) of the hippocampus mediates pattern separation by transforming overlapping or perceptually similar sensory inputs into distinct neural representations (O’Reilly and McClelland, 1994; Treves and Rolls, 1994; Leutgeb et al., 2007). The sparse activity of granule cells (GCs), wherein only a few GCs are active within a relevant time window with low firing rates, has been regarded to be essential for the computational function to accomplish pattern separation (O’Reilly and McClelland, 1994; Treves and Rolls, 1994; Rolls, 2013).

A hallmark of the DG is the continuous production of new neurons. During the period that newly generated GCs become fully mature to constitute a functionally homogenous neuronal population with developmentally born mature GCs(van Praag et al., 2002; Laplagne et al., 2006), the density of glutamatergic and GABAergic inputs and intrinsic excitability of GCs change enormously. Newly generated young GCs at age of 4-weeks display a low activation threshold due to high intrinsic excitability and enhanced excitation/inhibition balance, while mature GCs have a higher activation threshold due to strong inhibitory inputs and low intrinsic excitability (Schmidt-Hieber et al., 2004; Mongiat et al., 2009; Marin-Burgin et al., 2012; Dieni et al., 2013; Lopez-Rojas et al., 2016). However, how such different characteristics of young and mature GCs are relevant to DG functions is not fully understood. Most previous studies focused the role of adult-born young GCs in pattern separation (Clelland et al., 2009; Nakashiba et al., 2012), but it remains uncertain how the highly excitable young GCs can contribute to sparse and orthogonal dentate representation required for pattern separation function. The characteristic features of mature GCs, strong inhibitory inputs and low intrinsic excitability, may underlie sparse activity of GCs. Indeed, the hyperexcitability of GCs induced by selective degeneration of hilar mossy cells causing disinhibition led to impaired pattern separation (Jinde et al., 2012). However, whether low intrinsic excitability is critical for pattern separation has not been investigated. To directly test this possibility, identification of the mechanism specific to mature GCs’ low excitability is prerequisite.

Kv4.2 and Kv4.3 are expressed abundantly in the brain (Baldwin et al., 1991; Pak et al., 1991; Serodio et al., 1996), and encode A-type K^+^ channels in association with Kv channel interacting proteins (Rhodes et al., 2004) playing a key role in the regulation of neuronal excitability in various brain regions including the hippocampus (Hoffman et al., 1997; Locke and Nerbonne, 1997; Shibata et al., 2000; Ramakers and Storm, 2002). Kv4.1, which belongs to Kv4 gene subfamily, is known to have similar electrophysiological and pharmacological properties when expressed in heterologous expression system (Baldwin et al., 1991; Pak et al., 1991; Serodio et al., 1996), but its characteristics and roles in hippocampal neurons are largely unknown. In situ hybridization histochemistry showed that Kv4.1 signals are relatively strong in GCs compared to CA1 or CA3 pyramidal neurons (Serodio and Rudy, 1998), suggesting a possibility that Kv4.1 may play a specific role in regulating excitability of GCs.

In the present study, we demonstrate that Kv4.1 is expressed preferentially in mature GCs with a subcellular distribution pattern distinctive from Kv4.2. With its unique electrophysiological properties distinct from classical I_A_, Kv4.1 contributes to the low firing rates of GCs without affecting their passive electrical properties. Furthermore, we show that selective Kv4.1 depletion in the DG leads to the impairment in contextual discrimination task, suggesting that the low rate firing of mature GCs is a key component required for pattern separation.

## Materials and Methods

### Animals

All experiments were performed with C57BL/6 mice. Animals were housed 3–5 mice per cage and maintained under specific pathogen-free (SPF) conditions with food and water freely available. Electrophysiological analyses for the dentate gyrus (DG) granule cells (GCs) were performed in 4-to 8-week-old mice of both sexes, while experiments for CA1 pyramidal cells (PCs) were conducted using 3-to 5-week-old mice. Behavioral analyses were performed with 10-week-old male mice. The animal maintenance protocols and all experimental procedures were approved by the Institutional Animal Care and Use Committee (IACUC) at Seoul National University (SNU-090115-7, SNU-160825-1).

### Slice electrophysiology

Mice were sacrificed by decapitation after being anesthetized with isoflurane, and the whole brain was immediately removed from the skull, and chilled in artificial cerebrospinal fluid (aCSF) at 4 °C. Transverse hippocampal slices (350 μm thick) were prepared using a vibratome (VT1200S, Leica, Germany). Slices were incubated at 35 °C for 30 min and thereafter maintained at 32 °C until recording.

Hippocampal neurons were visualized using an upright microscope equipped with differential interference contrast (DIC) optics (BX51WI, Olympus, Japan). Electrophysiological recordings were made by the whole-cell patch clamp technique with EPC-8 or EPC-10 amplifier (HEKA, Lambrecht, Germany). Experiments were performed at 32 ± 1 °C. After membrane break-in, 2 ∼ 3 min wait time was given to stabilize neurons. Patch pipettes with a tip resistance of 3.5 – 4 MΩ were used. Series resistance (R_s_) after establishing whole-cell configuration was between 10 and 15 MΩ. Cells were discarded when the R_s_ changed by > 20% of baseline value. The internal pipette solution contained (in mM): 143 K-gluconate, 7 KCl, 15 HEPES, 4 MgATP, 0.3 NaGTP, 4 Na-ascorbate, and 0.1 EGTA, with the pH adjusted to 7.3 with KOH. For the antibody-blocking experiments, the Kv4.1 antibody was included in the pipette solution (0.3 µg/ml). The bath solution (or aCSF) for control experiments contained the followings (in mM): 125 NaCl, 25 NaHCO3, 2.5 KCl, 1.25 NaH2PO4, 2 CaCl2, 1 MgCl2, 20 glucose, 1.2 Na-pyruvate and 0.4 Na-ascorbate, pH 7.4 when saturated with carbogen (95% O2 and 5% CO2).

For current-clamp experiments to analyze neuronal excitability, 20 μM bicuculline and 10 μM CNQX were included in bath solutions to block synaptic inputs, and the following parameters were measured: (1) resting membrane potential, (2) input resistance (membrane potential changes at a given hyperpolarizing current input (−35 pA, 600 ms)), (3) F-I curve (firing frequencies (F) against the amplitude of injected currents (I)) (4) AP onset time (the delay from the start of the depolarized current injection to the beginning of the upstroke phase of the 1^st^ evoked AP), (5) AP half-width (measured as the width at 50% of the spike peak amplitude), (6) Overshoot (difference in voltage of AP peak amplitude from 0 mV), (7) AP threshold (the voltage at the point of deflection for d*V*/d*t* > 40 mV/ms). Membrane potentials are given without correction for liquid junction potentials.

For voltage-clamp experiments to analyze outward K^+^ currents, TTX (0.5 µM), CdCl2 (300 µM) and NiCl2 (500 µM) were additionally applied to block Na^+^ and Ca^2+^ channels, respectively. Outward K^+^ currents were evoked by 1-s voltage steps to potentials between –60 mV to +50 mV from a holding potential of –70 mV. After recording currents in the control condition, we added tetraethylammonium chloride (TEA) and 4-Aminopyridine (4-AP) sequentially for pharmacological dissection of K^+^ currents. The difference currents between outward K^+^ currents in control and those in 3 mM TEA were regarded as TEA-sensitive currents (ITEA), while the difference currents obtained after applying 5 mM 4-AP were regarded as 4-AP-sensitive currents (I4-AP).

For measurement of EPSC or EPSP in hippocampal neurons, an extracellular stimulation electrode was placed 100 – 150 μm away from the DG granule cell layer (GCL) or CA1 pyramidal cell layer, where stimulation was delivered to the medial perforant pathway (MPP) or Schaffer collateral (SC) pathway, respectively. EPSPs or EPSCs were evoked by 5 pulses at 50 Hz delivered by an isolation unit (ISO-Flex, AMPI, Israel) through monopolar electrodes (1.2 – 1.8 MΩ) filled with aCSF solution, and recorded in a whole-cell mode at a holding potential of –80 mV in mature GCs, and –63 mV in CA1-PCs. Stimulation intensity was adjusted such that EPSC amplitude was in the range between 150 pA and 250 pA. Picrotoxin (PTX, 100 µM) was included in the bath solution to block the GABAA receptors. All chemicals were obtained from Sigma-Aldrich (St. Louis, MO, USA), except CNQX, bicuculline, and TTX, which were from Abcam Biochemicals (Cambridge, UK).

### Immunofluorescence staining and western blotting

For the immunofluorescence staining, C57BL/6 mice were sacrificed at 3 or 8 weeks of age. AAV-injected mice were sacrificed at 12 weeks of age. Mice were anesthetized with isoflurane and perfused transcardially with a freshly prepared solution of 1X phosphate buffered saline (PBS) and 4% paraformaldehyde (PFA) for 10 – 15 min. Brains were removed and cut into 40 μm-thick sections using a vibratome (VT1200S, Leica, Germany), and post-fixed overnight at 4 °C by submersion in 4% PFA. After washing several times in 1X PBS with 0.3% Triton X-100 (PBS-T) for 5 min, sections were incubated 3 times in a blocking solution (2.5% donkey serum + 2.5% goat serum or 5% donkey serum in 0.3% PBS-T) for 1 h at room temperature (RT). The sections were then incubated overnight at 4 °C in blocking solutions containing the primary antibodies (anti-Kv4.1 antibody, APC-119, Alomone lab; anti-Kv4.2 antibody, APC-023, Alomone lab; anti-Kv4.3 antibody, APC-017, Alomone lab; anti-doublecortin (DCX) antibody, C-18, Santa Cruz biotechnology). After washing 5 times in 0.3% PBS-T for 5 min, sections were incubated with goat anti-rabbit Cy5 antibody (ab97077, Abcam), donkey anti-rabbit-Alexa488 (A21206, Invitrogen), and donkey anti-goat Alexa633 (A21082, Invitrogen) diluted in blocking solution for 1 h at RT. Sections were washed 5 times in 0.3% PBS-T for 5 min, and incubated with DAPI in PBS (Sigma-Aldrich) for 5 min. After rinsing with PBS, sections were mounted on glass slides using a fluorescent mounting medium (DakoCytomation, Cambridge, UK). The immunostained sections were imaged with a confocal laser scanning microscope (FV1200, Olympus, Japan) using a 40X oil-immersion objective or 60X water-immersion objective, and then processed using Fluoview (Olympus, Japan).

For western blotting, DG or CA1 region was isolated from slices under the dissecting microscope. Isolated tissues were homogenized with a glass homogenizer in TNE buffer (50 mM Tris-HCl, pH 8.0, 150 mM NaCl, and 2 mM EDTA) supplemented with protease inhibitor cocktails (Roche), and sonicated for 10 s. After adding SDS (0.5%) and Triton X-100 (1%), lysates were incubated for 30 min at 4 °C. Insoluble materials were removed by centrifugation at 20,000 × g for 15 min at 4 °C. Supernatants were mixed with 6× Laemmli buffer, separated by SDS-PAGE, and transferred to a PVDF membrane. Membranes were blocked in 5% skim milk in TBS for 1 h, and then probed with the relevant antibodies as indicated. Membranes were then incubated with peroxidase-conjugated secondary antibodies, and blots were detected with chemiluminescent reagents (Thermo Scientific).

### HEK293 cell electrophysiology and immunocytochemistry

HEK293 cells were cultured in Dulbecco’s modified Eagle’s medium (DMEM) supplemented with 10% (v/v) FBS and 1% (v/v) penicillin/streptomycin in a humidified incubator supplied with 5% CO2 at 37 °C. HEK293 cells were plated in a 12-well plate at a density of 1 × 10^5^ or 0.5 × 10^5^ cells per well for electrophysiology, and transfected with the Kv4 construct either alone (Kv4.1 or Kv4.2) or together with a GFP construct (Kv4.3) using Lipofectamine 2000 (Thermo Scientific) at a ratio of 1:6 (DNA/lipid). The GFP-tagged Kv4.1 (Cat# MG220056), GFP-tagged Kv4.2 (Cat# MG209597) and Myc-tagged Kv4.3 (Cat# MR221003) expression constructs were purchased from OriGene (Rockville, MD, USA).

Transfected HEK293 cells were maintained in an incubator for 1-2 days for the expression of Kv4.1, Kv4.2 or Kv4.3, and then transferred to a recording chamber where bath solution was perfused at 1 ml/min. The bath solution contained (in mM, 300 ± 10 mOsm): 143 NaCl, 5.4 KCl, 5 HEPES, 1.8 CaCl2, 0.5 MgCl2, 5 glucose with pH adjusted to 7.4 with NaOH. K^+^ currents were recorded by whole-cell patch clamp using a pipette solution of the following composition (in mM, 300 ± 10 mOsm): 110 K-aspartate, 30 KCl, 10 HEPES, 5 MgATP, 1 MgCl2, with the pH adjusted to 7.2 with KOH. Experiments were performed at room temperature (RT). For antibody-blocking experiments, patch pipettes were filled with the internal solution containing the Kv4.1 antibody (6 µg/ml). To measure K^+^ currents, we applied a 500 ms depolarizing voltage of +40 mV from a holding potential of −80 mV. Patch pipette resistance and series resistance (Rs) after establishing whole-cell configuration were 3-4 MΩ and 5-11 MΩ, respectively.

For immunocytochemistry, HEK293 cells were plated on poly-L-lysine (Sigma) coated coverslips in a 12-well plate and transfected with Kv4.1 or Kv4.2 cDNA. Cells were washed and fixed with 4% paraformaldehyde/4% sucrose in PBS for 15 min, and then permeabilized with 0.25% Triton X-100 for 5 min. Afterwards, they were blocked in PBS containing 1% normal goat serum for 1 h and incubated with primary antibody overnight at 4 °C. Coverslips were then incubated with Alexa 568-conjugated secondary antibody at RT for 1 h and mounted on slides. Images were captured using an Olympus FV 1000 laser scanning confocal microscope with a 63× objective.

### AAV production

Short hairpin RNA (shRNA) cassettes targeting murine Kv4.1 sequence (GCTGCCTTCTGGTATACCATT) or containing a non-targeting control sequence (TCGCATAGCGTATGCCGTT) were cloned under mU6 promoter of pAAV-U6-GFP vector (Cell Biolabs). AAV particles were produced in HEK 293T cells by co-transfecting pAAV-RC, pHelper, and rAAV vectors using calcium phosphate precipitation method. 72 hours after transfection, cells were lysed by three times of thaw/freeze cycles and the crude viral supernatant was obtained after a centrifugation at 12,000 X *g* for 30 min at 4 °C. After the viral supernatant was overlaid on iodixanol (OptiPrep™; Axis Shield) discontinuous density gradient (15/25/40/54%), AAV particles were collected at a 40% iodixanol density layer following a centrifugation at 350,000 X *g* for 90 min at 18 °C. AAV particles were dialyzed and concentrated using Amicon Ultra-15 centrifugal filter with Ultracel-100 membrane (Millipore). Aliquoted viral stock (∼5 × 10^12^ GC/ml) was stored at −80 °C before use.

### Stereotaxic surgery

5-week-old mice were anaesthetized and stereotaxically injected with high titers (∼5 × 10^12^ GC/ml) of AAVs (1 μl). The injection was performed through glass capillary at 4 sites at the following coordinates relative to bregma (mm): AP:-2, ML: ±1.6, DV:-2.5; or AP:-3, ML: ±2.6, DV:-3.2. After infusion, the glass capillary was left in place for an additional 5 min to ensure full virus diffusion. After surgery, mice were treated with antibiotics daily for two weeks, and their health was monitored every day.

### Behavior analysis

Contextual fear conditioning and discrimination task were performed three weeks after surgery. Mice were trained to discriminate between two similar contexts, A and B, through repeated experience in each context. Context A (conditioning context) was a chamber (18 cm wide x 18 cm long x 30 cm high; H10-11M-TC; Coulbourn Instruments 5583, PA 18052, USA) consisting of a metal grid floor, aluminum side walls, and a clear Plexiglass front door and back wall. Context A was indirectly illuminated with a 12 W light bulb. The features of Context B (safe context) were the same as Context A, except for a unique scent (1% acetic acid), dimmer light (50% of A), and a sloped floor by 15° angle. Each context was cleaned with 70% ethanol before the animals were placed. On the first 3 days (contextual fear acquisition), mice were placed in Context A for 3 min for exploring the environment, and then received a single foot shock (0.75 mA, for 2 s). Mice were returned to their home cage 1 min after the shock. On day 4 – 5, mice of each genotype were divided into two groups; one group visited Context A on Day 4 and Context B on Day 5, while the other group visited the Context B on Day 4 and Context A on Day 5. On day 4 – 5 (generalization), neither group received a shock in Context A and B, and freezing level was measured for 5 min only in Context A. We defined freezing behavior as behavioral immobility except for respiration movement (McNaughton and Nadel, 1990). We observed video image for 2-s bouts every 10 s (18 or 30 observation bouts for 3 min or 5 min recording time) and counted the number of 2-s bouts during which the mouse displayed freezing behavior (referred to as the freezing score). The percentage of freezing was calculated by dividing the freezing score with the total number of observation bouts (18 or 30). Mice were subsequently trained to discriminate these two contexts by visiting the two contexts daily for 7 days (from day 6 to 12, discrimination task). Mice always received a footshock (2 s) 3 min after being placed in Context A but not B. Discrimination ratios were calculated according to FA / (FA + FB), where FA and FB are freezing scores in Contexts A and B, respectively. All experiments and analyses were performed blind to the mice genotype.

Open field exploration test (OFT) was performed in white plastic boxes (40 * 40 * 40 cm) in a dimly lit room. The open field was divided into a center zone (center, 20 * 20 cm) and an outer field. Individual mice were placed in the center zone, and the path of the animal was recorded with a video camera. Total distance traveled was measured for the entire period (20 min) and time spent in the center zone was analyzed for initial 5 min.

Elevated plus maze (EPM) test was performed using a plus-shaped apparatus consists of two open arms and two closed arms. Each arm was 65 * 6.25 cm in size, and the maze was elevated 50 cm above the ground. Mice were individually placed on the center area, facing an open arm, and allowed to freely explore the apparatus for 5 min. We analyzed the total distance traveled for 5 min and time spent in each arm. OFT and EPM were analyzed using EthoVision XT (Noldus, Netherlands)

Novel object recognition tests were conducted in the same open field boxes used for the OFT. Two identical objects were placed in the box during the familiarization phase, and one of the objects was replaced with a similar new object during the discrimination test. The interval between the familiarization phase and the discrimination test was 24 h. In each session, mice were allowed to explore freely the objects, and exploratory activity was recorded for 10 min. Discrimination index was calculated as the time spent exploring a similar novel object divided by the total time spent exploring both the novel and the familiar objects.

### Statistical analysis

Data were analyzed with Igor Pro (Version 6; Wavemetrics, Lake Oswego, CA) and Origin (Version 8; Microcal, Northampton, MA). All results are presented as mean ± SEM with the number of cells or mice (n) used in each experiment. Statistical significance was evaluated using Student’s t-test, and the level of significance was indicated by the number of marks (*, P < 0.05; **, P < 0.01; ***, P < 0.001). P > 0.05 was regarded as not significantly different (N.S.). Comparison between multifactorial statistical data was made using the two-way analysis of variance (ANOVA). Differences in time-dependent changes of behavioral parameters between the two genotypes were evaluated using two-way repeated measures ANOVA.

## Results

### Preferential expression of Kv4.1 in the DG

Among Kv4 family channels that mediate IA, Kv4.2 is highly expressed in principal neurons throughout the hippocampus and known to play crucial roles in regulating intrinsic excitability and synaptic plasticity (Kim et al., 2005; Chen et al., 2006; Kim et al., 2007). However, little is known about the distribution and functions of Kv4.1. We first investigated the distribution and expression of Kv4.1 using the Kv4.1-specific antibody in comparison with Kv4.2 in different subregions of the hippocampus obtained from adult mouse (8-week-old). Antibody specificity was confirmed using HEK293 cells expressing Kv4.1 or Kv4.2 isoform alone (Fig. 1A). Immunohistochemistry and western blot analyses showed that both Kv4.1 and Kv4.2 were expressed in the hippocampus, but their regional and subcellular distributions were different. Specifically, Kv4.1 signal was evident in the DG and CA3, but it was very weak in CA1, while Kv4.2 signal was strong both in the DG and CA1, but weak in CA3 (Fig. 1B). Moreover, the two isoforms showed distinct patterns of subcellular localization. Kv4.1 signals were localized mainly in the principal cell layer of the hippocampus, whereas Kv4.2 signals were localized mainly in dendritic regions (Fig. 1B). Average fluorescence intensities for Kv4.1 and Kv4.2 obtained from the DG and CA1 subregions and western blot analysis confirmed these observations (Figs. 1C,D).

**Figure 1.**
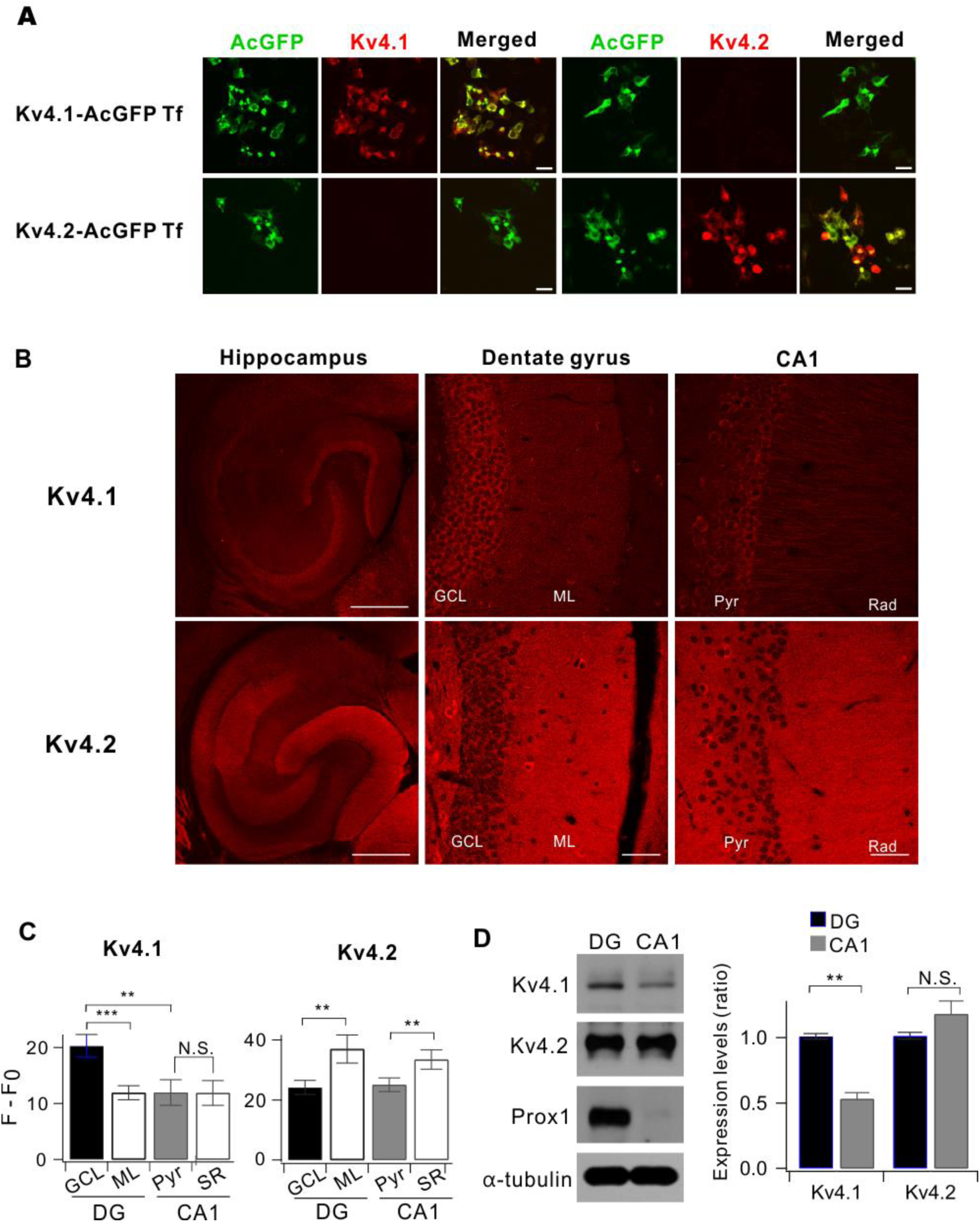
Comparison between Kv4.1 and Kv4.2 expression in the hippocampus. (A) Both Kv4.1 and Kv4.2 antibodies selectively recognize the channel in HEK293 cells. HEK293 cells were transfected with either AcGFP-tagged Kv4.1 cDNA (Kv4.1-AcGFP Tf) or Kv4.2 (Kv4.2-AcGFP Tf). Forty-eight hours after transfection, the cells were stained using Kv4.1 or Kv4.2 antibodies. Scale bar, 20 μm. (B) Representative fluorescence immunostaining images show that Kv4.1 (top) is highly expressed in DG while Kv4.2 (bottom) is broadly expressed in the hippocampus of 8-week-old mice. Scale bars, 500 (left) and 50 (middle and right) µm. (C) Summary bar graphs of the relative fluorescence intensity (F-F_0_) shown in the DG and CA1 for Kv4.1 (n = 10, left) and Kv4.2 (n = 5, right). (Kv4.1; GCL, 20.3 ± 2.0; ML, 11.9 ± 1.3; Pyr, 12.0 ± 2.3; Rad, 11.9 ± 2.2; GCL vs. ML, p < 0.005; GCL vs. Pyr, p < 0.01; Pyr vs. Rad, p = 0.75; Kv4.2; GCL, 24.2 ± 2.3; ML, 39.1 ± 5.8; Pyr, 25.1 ± 2.3; Rad, 33.5 ± 3.3; GCL vs. ML, p < 0.01; GCL vs. Pyr, p = 0.6; Pyr vs. Rad, *P* < 0.01, paired t-test). (D) Western blot analysis for Kv4.1 and Kv4.2 in DG and CA1 of 8-week-old mice. Right, DG exhibits higher Kv4.1 expression level compared to CA1; 1.00 ± 0.02 in DG, 0.53 ± 0.05 in CA1, n = 8; *P* < 0.0001. Paired t-test. Prospero-related homeobox 1 (Prox) and α-tubulin used as a marker for DG and loading control, respectively.

### Kv4.1 inhibition selectively increases the firing frequency of low-frequency-firing GCs

To investigate the functional significance of Kv4.1 expression in GCs, we generated an adeno-associated virus (AAV) vector to stably express a short hairpin RNA (shRNA) against the Kv4.1 transcript (shKv4.1). In contrast to non-targeting shRNA sequence (shCtrl), infection of shKv4.1 significantly reduced Kv4.1 expression in cultured hippocampal neurons, whereas the expression levels of both Kv4.2 and Kv4.3 were not affected by the shKv4.1 infection, confirming the specificity of shKv4.1 (Fig. 2A). We then examined the effect of Kv4.1 knockdown on the electrical properties of GCs 2 weeks after stereotaxic injection of shKv4.1 or shCtrl into the DG. To avoid possible sources of confounding factors such as the neural progenitor cells and newborn immature neurons, we chose GCs in the outer granular cell layer (GCL) where mature GCs are located (Kerloch et al., 2018). Resting membrane potential (RMP) and input resistance (Rin) in shKv4.1-infected GCs (−81.8 ± 1.05 mV and 146.1 ± 8.9 MΩ, n = 11) and those in shCtrl-infected GCs (−82.4 ± 2.0 mV and 155.1 ± 26.9 MΩ, n = 9) showed no significant difference between groups (Fig. 2B), and the values were consistent with those reported previously for mature GCs (Schmidt-Hieber et al., 2004; Dieni et al., 2013). When we induced repetitive APs using a depolarizing current injection, however, the firing rate was significantly higher in shKv4.1-infected GCs than in shCtrl-infected GCs (Fig. 2Ca). The relationships between the firing frequency and input current (F-I curves) shifted upward in shKv4.1-infected GCs with an increase in slope, indicating the increased gain of input-output relationship (I-O gain) (Fig. 2Cb). I-O gain calculated from the frequency increment between 100 and 200 pA input current was significantly higher in shKv4.1-infected GCs (20.2 ± 1.7, n = 9) than in shCtrl-infected GCs (6.6 ± 1.4, n = 11). These results suggest that Kv4.1 channels do not affect the passive membrane properties, but contribute to the low firing rate of GCs.

**Figure 2.**
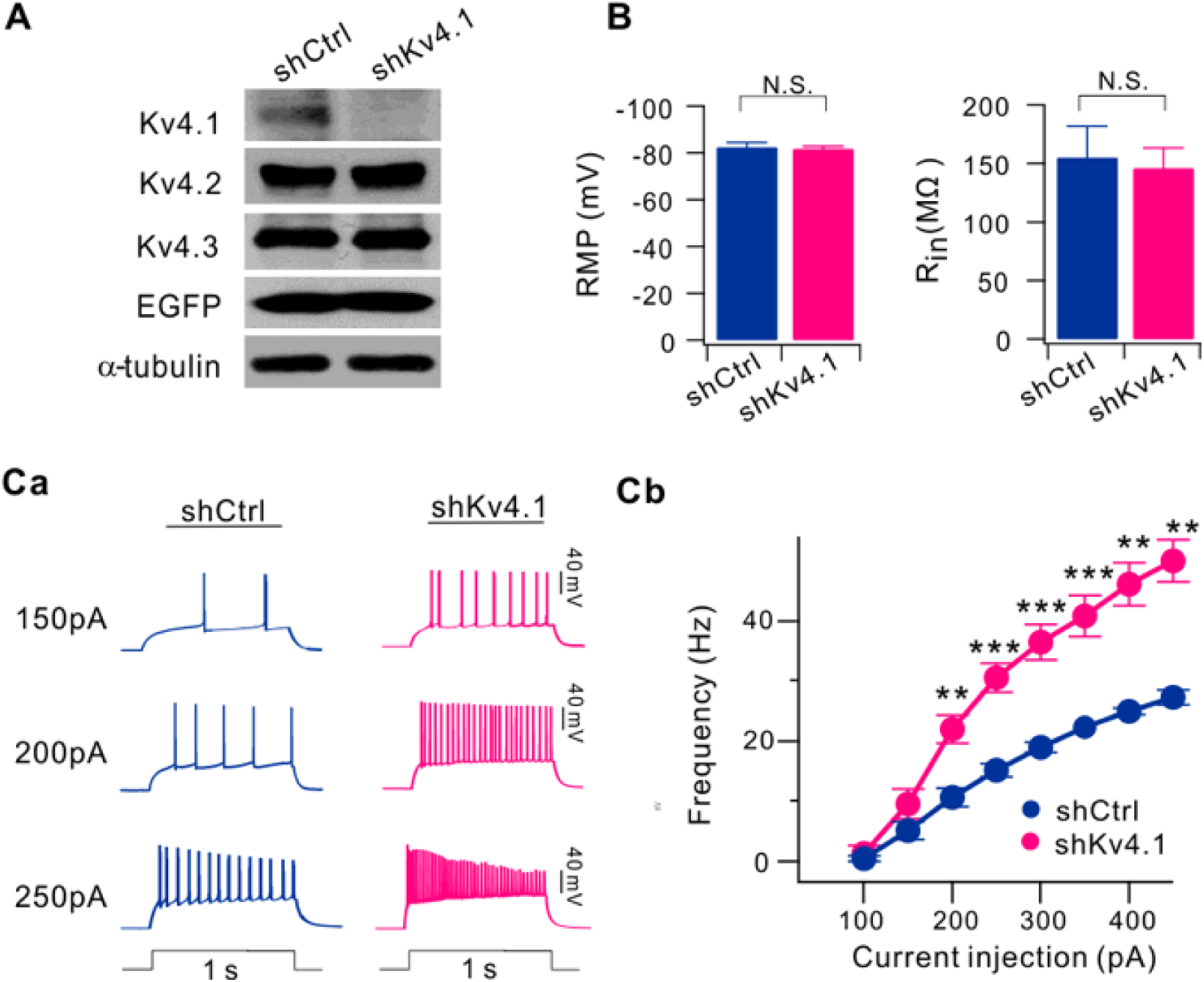
Knockdown of Kv4.1 increases firing frequency in GCs. (A) Specificity and efficiency of Kv4.1 knockdown in cultured hippocampal neurons. Cultured hippocampal neurons were infected by AAV-shCtrl or AAV-shKv4.1 at DIV8 for 7 days. At DIV15, endogenous expression levels of Kv4.1 were analyzed by western blotting. (B) Knockdown of Kv4.1 had no effect on both RMP and R_in_ in GCs. (Ca) Representative traces of the AP train recorded from the GCs in brain slices from shCtrl- and shKv4.1-injected mice in response to 150, 200 and 250 pA current injection (1-s duration). (Cb) Averaged F-I curve in GCs infected with AAV-shCtrl (dark blue, n = 11) or AAV-shKv4.1 (pink, n = 9).

Electrical properties of GCs are not homogenous, since they change during maturation, in a way that Rin decreases and RMP hyperpolarizes with time (Schmidt-Hieber et al., 2004; Mongiat et al., 2009; Dieni et al., 2013). To further study the role of Kv4.1 in firing frequency regulation, we first examined the heterogeneity of firing properties among GCs with Rin between 100 and 600 MΩ where depolarizing current injection evoked repetitive firing faithfully. Cells with Rin in this range exhibited typical morphology of GCs (Fig. 3A), and GCs with lower Rin were found in the outer granule cell layer (GCL), while GCs with higher Rin were mostly found in the inner GCL. The relative positions of the cells shown in Fig. 3A in the GCL were 0.19, 0.3, and 0.69 when the GCL was scaled from 0 to 1 from the border of molecular layer to the hilus. RMP varied widely between −50 to −90 mV, in a way that RMP hyperpolarization occurred steeply as Rin decreased lower than 300 MΩ (Fig. 3B). The firing frequency of repetitive APs induced by long depolarization (1-s duration) also varied greatly ranging from a few to more than 50. It was interesting to note that the distribution of firing frequency was not as continuous as Rin or RMP (Fig. 3B), but can be categorized into two distinctive groups (Figs. 3C). One group of GCs was hardly activated in response to a depolarizing pulse of 100 pA, and the number of APs induced by 200 pA current injection was less than 12, while the other group of GCs was much more excitable, and the number of APs induced by 200 pA current injection was far above20. We therefore referred to the former group as the low-frequency-firing GCs (LF-GCs) and the latter group as the high-frequency-firing GCs (HF-GCs). The relationship between firing frequency and Rin or RMP showed a tendency that firing frequency increased with Rin and RMP depolarization. Most LF-GCs had Rin smaller than 200 MΩ and RMP lower than −75 mV (Figs. 3Cb, Cc, blue symbols). The mean Rin value for LF-GCs (153.6 ± 6.6 MΩ n = 20) was significantly smaller than the value for HF-GCs (376.8 ± 21.3 MΩ n = 27, P < 0.001), and the mean RMP value for LF-GCs (−82.9 ± 1.2 mV, n = 20) was significantly lower than that for HF-GCs (−64.1 ± 1.8 mV, n = 27, *P* < 0.001). In contrast, the properties of single AP, such as AP threshold, overshoot, and AP width, were fairly constant in GCs with Rin in this range, and there was no significant difference between LF-GCs and HF-GCs (Fig. 3D). These results suggest that ion channels required for single AP generation mature before Rin reaches about 600 MΩ while ion channels responsible for RMP hyperpolarization and lowering frequency mature at a late stage.

**Figure 3.**
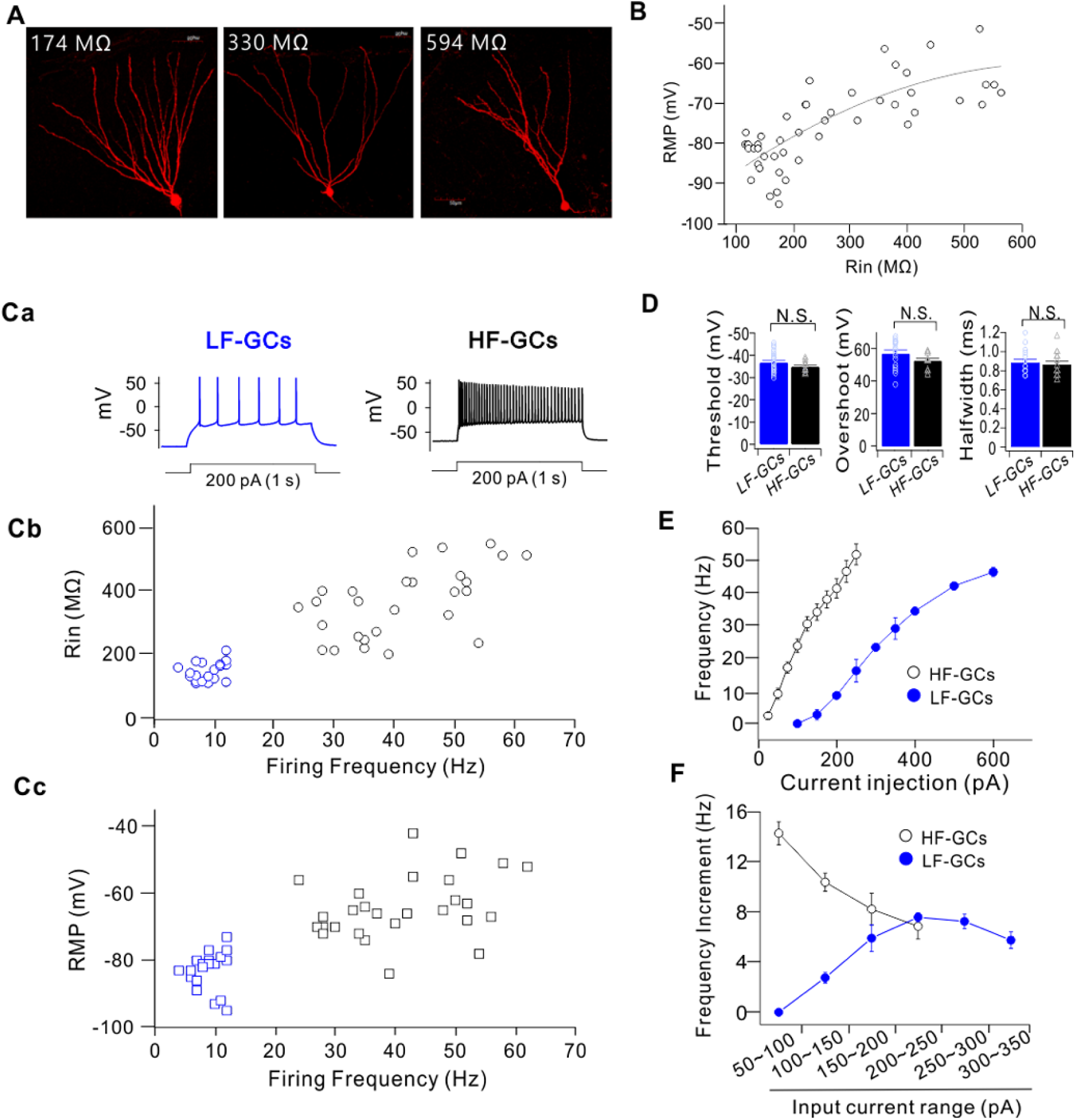
GCs cluster into two distinct populations based on the firing frequency. (A) Sample images of GCs with different R_in_ values. (B) Plots of RMP values against R_in_. Data obtained from 47 cells are fitted to a polynomial function: RMP (mV) = −97.6 + 0.11 R_in_ + 0.000087 R_in2_. (C) Firing frequency of the repetitive APs evoked by a depolarizing current injection showed two distinct groups. Representative traces for APs of low-frequency (LF)- and high-frequency (HF)-GCs in response to 200 pA (1-s duration) current injection (Ca). Plots for R_in_ (Cb) and RMP (Cc) against firing frequency at 200 pA showed that firing frequency decreased with a decrease in R_in_ and RMP, reaching the lowest level at R_in_ below 200 MΩ and at RMP below −75 mV. (D) AP threshold, overshoot, and AP width were not significantly different between LF-GCs and HF-GCs. (E) The firing frequency plotted against the injected currents (F-I curve) in LF-(filled blue circle, n = 12) and HF-(open black circle, n = 13) GCs. HF-GCs fired more action potentials in response to smaller current injection than LF-GCs. (F) Increment in firing frequency for given increases in input currents in LF-(filled blue circle) and HF-(open black circle) GCs.

The difference in firing frequencies between LF-GCs and HF-GCs was further analyzed in F-I curves. The minimum current for activation was much larger, resulting in a rightward shift in the F-I curve in LF-GCs (Fig. 3E). Furthermore, the slope of the F-I curve was less steep in LF-GCs, indicating that the I-O gain was lower. I-O gain was analyzed by comparing the frequency increment for a given increase in input current, showing that the input current range where the I-O gain was maximum is different between LF-GCs and HF-GCs and that the maximum gain was significantly larger in HF-GCs (−7.58 ± 0.35, n = 12, between 200 and 250 pA for LF-GCs; −14.29 ± 0.95, n = 13, between 50 and 100 pA for HF-GCs, *P* < 0.001, Fig. 3F).

We then investigated whether Kv4.1 may have different roles in LF-GCs and HF-GCs. To this end, we used a specific antibody to Kv4.1 for the functional blockade of Kv4.1 channels. Application of the Kv4.1 antibody through a patch pipette robustly inhibited the Kv4.1 currents but not the Kv4.2 currents or Kv4.3 currents overexpressed in HEK293 cells (Figs. 4A, B), confirming the specificity of the Kv4.1 antibody. A typical example for effects of Kv4.1 antibody perfusion in LF-GCs was shown in Fig. 4C.

**Figure 4.**
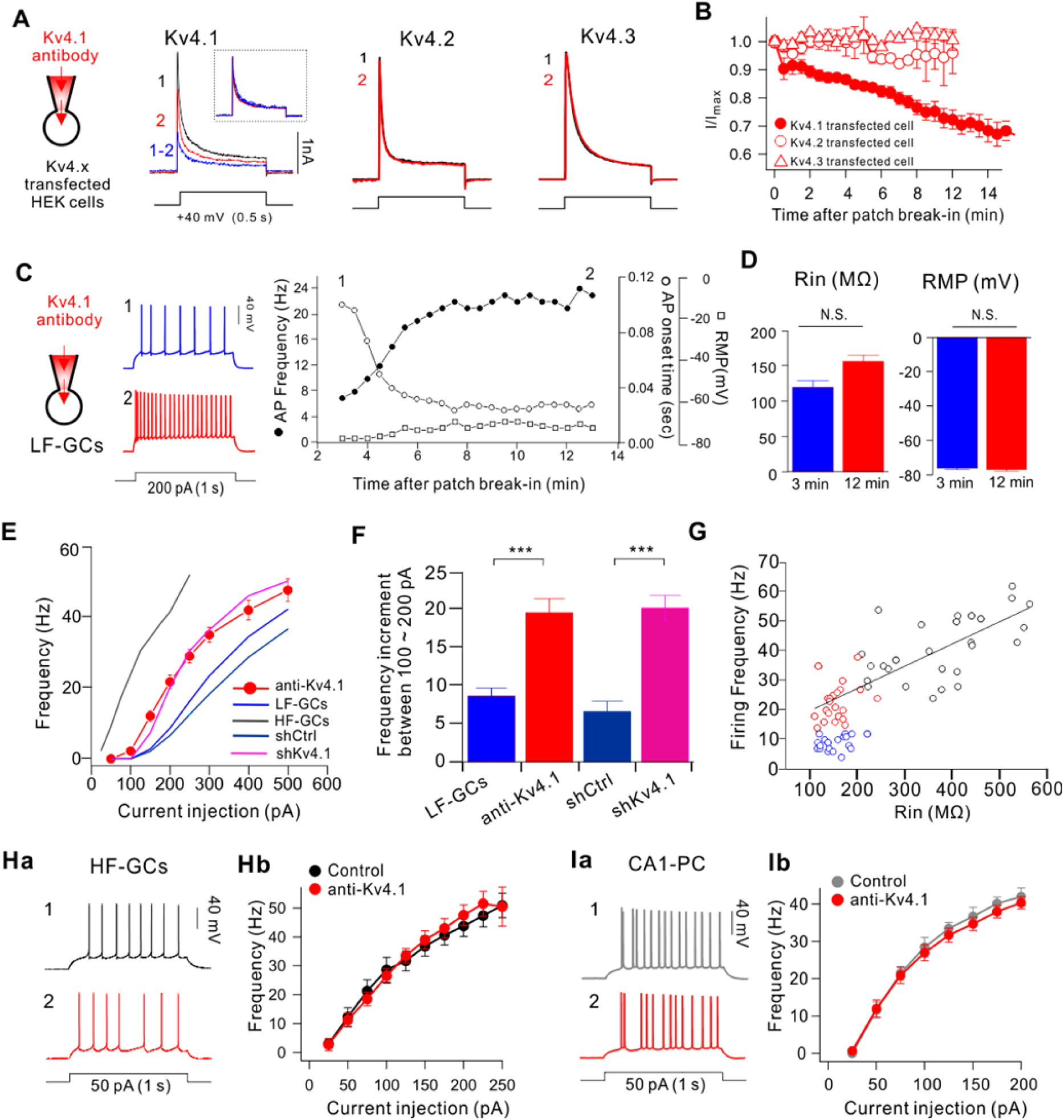
Kv4.1 inhibition increases firing frequency selectively in LF-GCs. (A) The specificity of Kv4.1 antibody for the Kv4.1-mediated current was determined in HEK293 cells expressing Kv4.1, Kv4.2 or Kv4.3. Outward currents evoked by depolarizing pulse to +40 mV for 1 s in Kv4.1, Kv4.2 or Kv4.3-expressing HEK293 cells measured just after patch break-in (1, black) and more than 12 min after intracellular application of the Kv4.1 antibody (2, red). Inset, K+ currents measured before (black) and after (red) inhibition of Kv4.1 and subtracted current (blue) in the Kv4.1-expressing cell were normalized to the peak amplitude of current. (B) Summary graph of inhibition of I/I_max_ by Kv4.1 antibody in HEK293 cells expressing Kv4.1 (closed circle), Kv4.2 (open circle), or Kv4.3 (open triangles). Values indicate mean ± SEM. Error bars indicate SEM. (C-H) Effects of Kv4.1 antibody on intrinsic excitability in LF-GCs (C-G) and HF-GCs (H). (C) Example traces (left) of APs at the time points (1, 2) indicated on the right graph. Right, Graphs showing the time course of the alterations in RMP (open squares), firing frequency (filled circles) and 1^st^ AP onset time to 1-s current injection (open circles) during Kv4.1 antibody perfusion in LF-GCs. (D) Mean values for Rin and RMP (n = 6) measured within 3 min after patch break-in and after the changes in firing frequency induced by Kv4.1 antibody reached steady state (12 min). (E) The F-I curve measured from LF-GCs in the presence of Kv4.1 antibody in the pipette solution (red closed circles, n = 10). The same data shown in Fig. 2Cb (dark blue, shKv4.1; pink, shCtrl) and Fig. 3E (blue, LF-GCs) are superimposed to compare with the F-I curve obtained in the presence of Kv4.1 antibody. (F) I-O gain between 100 and 200 pA input currents obtained from each F-I curve shown in Fig. 4E. Inceases in I-O gain by Kv4.1 inhibition were significant (*P* < 0.001). (G) Firing frequency of GCs under control condition (blue circles for LF-GCs; black circles for HF-GCs) and Kv4.1-depleted conditions using shKv4.1 or Kv4.1 antibody (red circles) was plotted against Rin. The firing frequency of HF-GCs (black circles) and Kv4.1-depleted LF-GCs (red circles) showed a linear relationship with R_in_, while the firing frequency of LF-GCs (blue circles) is lower. (H) The Kv4.1 antibody had no effects on intrinsic excitability in HF-GCs. Representative traces of AP trains obtained from HF-GCs recorded right after patch break-in (1, black) and 15 min after intracellular application of Kv4.1 antibody (2, red). (Hb) Inhibition of Kv4.1 had no effect on the firing frequency of HF-GCs (n = 4). (Ia) Representative traces from CA1-PCs recorded right after patch break-in (1, grey) and 15 min after intracellular application of Kv4.1 antibody (2, red). (Ib) Firing frequency was plotted against the injecting currents (1-s) with or without Kv4.1 antibody in the pipette for CA1-PCs (n = 7).

After patch break-in, AP firing in response to 200 pA current injection increased gradually (filled circles, Fig. 4C) in association with a shortening of the 1^st^ AP onset time (open circles, Fig. 4C), while RMP was not significantly changed (open squares, Fig. 4C). When mean firing frequency (8.6 ± 0.8 Hz) and the 1^st^ AP onset time (0.122 ± 0.008 s; n = 6) were obtained from the responses to the first stimulus, the values were significantly different from those obtained after changes reached steady state (17.0 ± 2.4 Hz and 0.063 ± 0.008 s, n = 6). The mean Rin value measured after the effect of Kv4.1 antibody reached steady state was slightly larger than the value measured just after patch break-in, but the difference was not statistically significant (156.7 ± 9.2 MΩ *vs* 119.7 ± 9.8 MΩ, n = 6, *P* > 0.01). Effect of Kv4.1 inhibition in LF-GCs was also confirmed by comparing F-I curves obtained from Kv4.1 antibody-perfused GCs (red circles, Fig. 4E) with those from control GCs shown in Fig. 3E. There was no significant shift in threshold current, but the slope increased markedly in Kv4.1 antibody-perfused GCs compared to the slope in control LF-GCs (blue line, Fig. 4E), and became comparable to the slope in HF-GCs (grey line, Fig. 4E).. We also superimposed F-I curves obtained in Fig. 2Cb, showing a similar increase in slope in shKv4.1-infected GCs (pink line, Fig. 4E) compared to shCtrl (dark blue line, Fig. 4E). I-O gain obtained between 100 and 200 pA input current in the presence of Kv4.1 antibody (19.6 ± 1.9, n = 10) was significantly higher than that in control LF-GCs (8.6 ± 1.1, n = 12, *P* < 0.001, Fig. 4F), and comparable to the value in shKv4.1-infected GCs (Fig. 4F) or I-O gain between 0 and 100 pA input current in HF-GCs (23.9 ± 2.2, n = 13).

To further understand the role of Kv4.1 in firing frequency regulation in GCs, we pooled all data obtained under different experimental conditions and plotted the firing frequencies of individual GCs induced by 200 pA injection against their Rin. Data obtained from HF-GCs under control conditions (black circles, Fig. 4G) together with data obtained under Kv4.1-inhibited conditions with shKv4.1 or Kv4.1 antibody (red circles, Fig. 4G) represent the relationship between Rin and firing frequency in the absence of Kv4.1, showing that firing frequency decreases with the decrease of Rin. However, data points for LF-GCs under control condition (blue circles, Fig. 4G) were below this relationship, suggesting that the low frequency of LF-GCs cannot be fully explained by low Rin, but Kv4.1 provides an additional outward shunting conductance upon depolarization to further lower firing frequency. In contrast, the firing frequency of HF-GCs was unaffected by Kv4.1 antibody (Figs. 4Ha, b). We also confirmed that the firing frequency of CA1-PCs was unaffected by Kv4.1 antibody (Figs. 4Ia, b), which is consistent with the weak expression of Kv4.1 in CA1-PCs (Fig. 1). These results suggest that Kv4.1 selectively contributes to frequency regulation in LF-GCs.

### Distinct biophysical properties of Kv4.1 currents underlie its function as a frequency regulator

The selective role of Kv4.1 in LF-GCs suggests a higher amplitude of Kv4.1 currents in LF-GCs. To verify this possibility, we tested the effects of Kv4.1 antibody on outward K^+^ currents evoked by a depolarizing voltage step from a holding potential of −70 mV to +30 mV (1 s duration). In voltage clamp experiments, we regarded GCs with Rin smaller than 200 MΩ as LF-GCs, while GCs with Rin ranging from 300 to 600 MΩ as HF-GCs, since LF-GCs and HF-GCs can be also distinguished with respective to Rin value of around 200 MΩ (Fig. 3Cb). We monitored the changes in peak outward current amplitudes (Ipeak) during intracellular dialysis of GCs with internal solution containing Kv4.1 antibody (0.3 µg/ml), and found that Ipeak decreased gradually to a significant extent, reaching a steady state usually within 15 min in LF-GCs (Figs. 5A, D). In contrast, the effects of Kv4.1 antibody on Ipeak were negligible in HF-GCs (Figs. 5A, D), LF-GCs from shKv4.1 transfected mouse (Figs. 5B, D), and CA1-PCs (Figs. 5C, D), where Kv4.1 had no significant role in regulating firing frequency (Figs. 4H, I). These results suggest that Kv4.1 significantly contributes to outward K^+^ current only in LF-GCs, leading to the selective role of Kv4.1 in LF-GCs.

**Figure 5.**
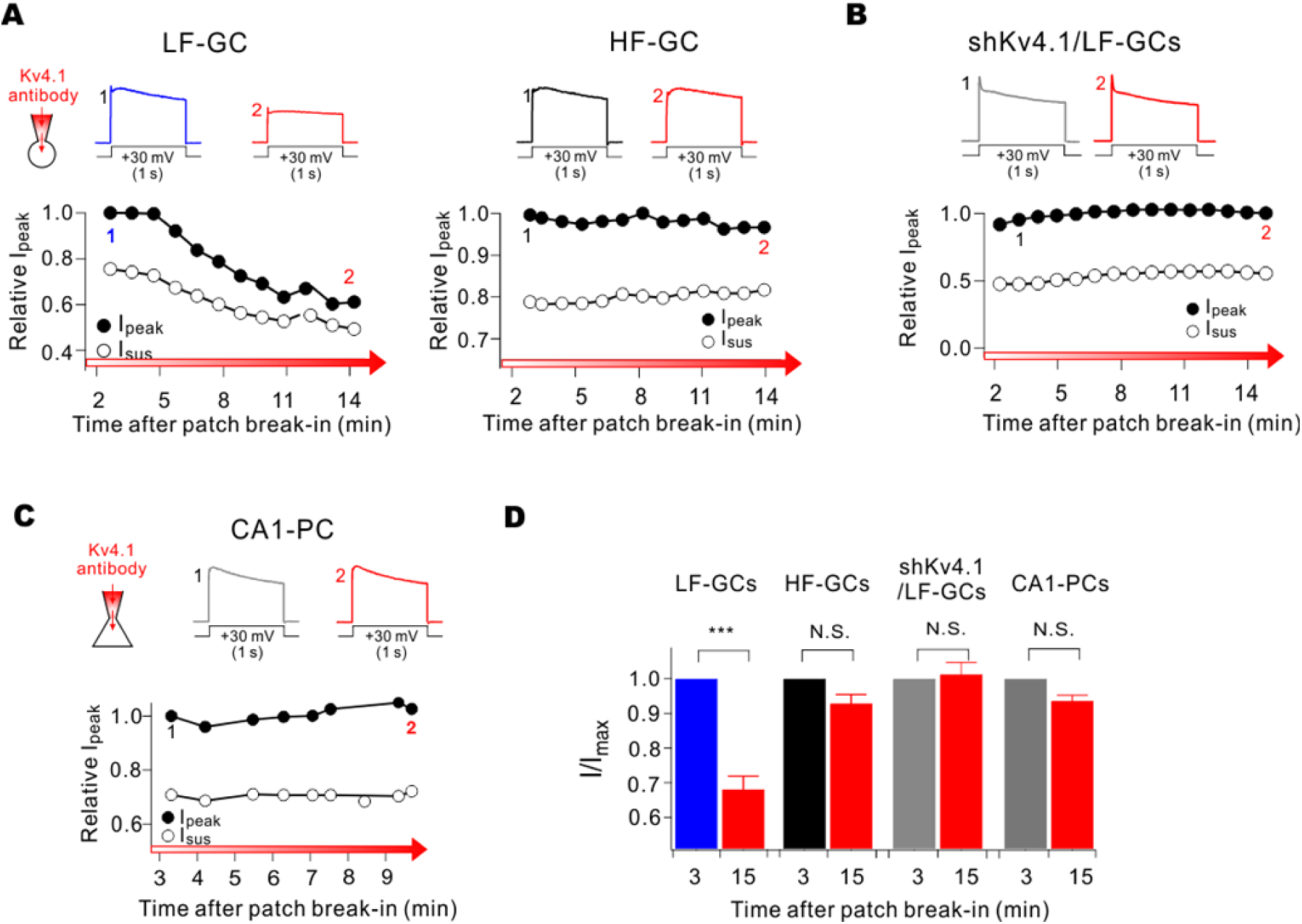
Kv4.1 inhibition reduces K^+^ current selectively in LF-GCs. (A-C) Effects of Kv4.1 antibody on outward K^+^ currents in LF- and HF-GCs (A), LF-GCs infected by shKv4.1 (B), and CA1-PCs (C). (top) Representative current traces recorded shortly after patch break-in usually within 3 min (marked with 1) and those recorded after 10 – 15 min (marked with 2). A significant decrease in outward K^+^ currents by Kv4.1 antibody was observed only in LF-GCs. Voltage protocol was depicted below current traces. (bottom) The current amplitude measured at the peak (I_peak_) normalized to the maximum amplitude during recording period were plotted against the time after patch break-in. (D) I_peak_ obtained at 15 min after patch break-in normalized to that obtained at 3 min after patch break-in were pooled and plotted as bars with SEM. LF-GCs (n = 15), HF-GCs (n = 8), shKv4.1-infected LF-GCs (n = 7), CA1-PCs (n = 9). Paired t-test. \*\*\**P* < 0.001.

To further understand the mechanism underlying the role of Kv4.1 as a frequency regulator in LF-GCs, we analyzed the electrophysiological characteristics of Kv4.1 currents. To this end, we first analyzed the difference in outward K^+^ currents between LF-GCs and HF-GCs. Whole-cell K^+^ currents evoked by applying depolarization pulses in a 10-mV step from the holding potnetial of −70 mV were dissected into TEA-sensitive currents (I_TEA_) and 4-AP-sensitive currents (I4-AP) using 3 mM TEA and 5 mM 4-AP, respectively. Average current traces (n = 9 for LF-GCs, n = 4 for HF-GCs) showed that the amplitude of ITEA was larger in LF-GCs compared to HF-GCs (Fig. 6A), but there was no significant difference when the amplitude was normalized to the cell capacitance (Fig. 6B), indicating that the density of ITEA was not significantly changed with the decrease in Rin in this range. With respect to I_4-AP_, not only the current amplitude (Fig. 6C) but also the current density normalized to the cell capacitance (Fig. 6Da) was significantly larger in LF-GCs compared to those of HF-GCs. Furthermore, there was a distinctive feature in I4-AP between LF- and HF-GCs. I4-AP in HF-GCs showed rapidly inactivating kinetics and the current amplitude measure at the end of the pulse (Iss) was negligible, whereas a considerable proportion of I4-AP in LF-GCs was not fully inactivated even after 1 s (Iss/Ipeak at +30 mV, 0.24 ± 0.02, n = 9, Fig. 6Db). The difference in I4-AP between LF- and HF-GCs may be attributable to the contribution of Kv4.1 currents in LF-GCs. Kv4.1 currents are known to have similar electrophysiological characteristics with Kv4.2 or Kv4.3 currents when studied in heterologous expression systems (Nakamura et al., 2001), but its kinetics in native neurons has not been well studied. Different kinetics of I_4-AP_ in LF- mand HF-GCs suggest that Kv4.1 current may have different kinetics from typical A-type K^+^ currents (IA). We obtained Kv4.1 currents (IKv4.1) from LF-GCs in the presence of 3 mM TEA to inhibit delayed rectifier K^+^ currents by subtracting the currents measured 15 min after patch break-in in the presence of Kv4.1 antibody from currents recorded immediately after break-in (Fig. 6E). Subtraction results showed that IKv4.1 inactivated slowly, and thus a significant proportion of currents remained at the end of the 1-s depolarization (red, Fig. 6E). When 5 mM 4-AP was applied after IKv4.1 was completely inhibited by the Kv4.1 antibody, there was further reduction in outward currents (Fig. 6F). The 4-AP-sensitive currents obtained by subtraction showed a rapid voltage-dependent activation followed by rapid inactivation, the characteristics of which are consistent with the kinetics known for classical A-type K^+^ currents.

**Figure 6.**
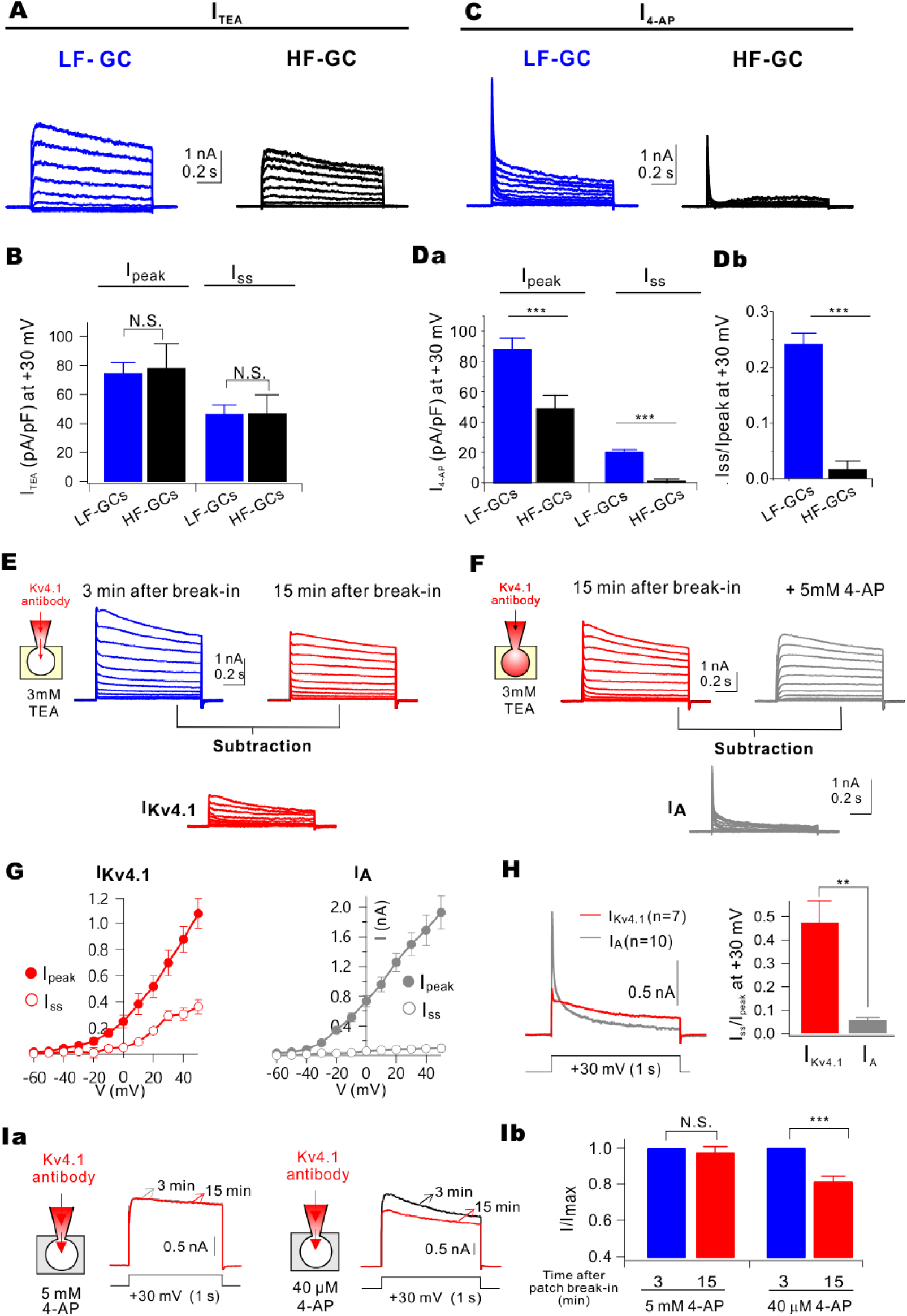
Kv4.1 contributes to slowly inactivating sustained outward currents in GCs. (A) Average current traces for TEA-sensitive currents (I_TEA_) in LF-GCs (blue, n = 9) and HF-GCs (black, n = 4). (B) Current density (pA/pF) of I_peak_ and I_sus_ at +30 mV for I_TEA_ was not different in LF-GCs (blue) and HF-GCs (black). (C) Average current traces for 4-AP-sensitive currents (I_4-AP_) in LF-GCs (blue, n = 9) and HF-GCs (black, n = 4). (Da) I_4-AP_ density in LF-GCs was significantly larger than that in HF-GCs (I_peak_, *P* < 0.001; I_sus_, *P* < 0.001). (Db) The ratio of I_ss_ to I_peak_ (I_ss_/I_peak_) for I_Kv4.1_ was significantly enhanced in LF-GCs (*P* < 0.001). (E) Whole-cell, voltage-gated K^_+_^ currents in LF-GCs at the beginning of the recording (blue) and after 15 min dialysis of Kv4.1 antibody (red). The difference traces shown below (red) represent currents mediated by Kv4.1 channel. (F) Records showing subsequent effects of 5 mM 4-AP after the maximal effect of Kv4.1 antibody in LF-GCs. The difference traces (grey) are defined as A-type K^+^ current (I_A_). (G) Averaged current-voltage (I-V) relationships of difference traces at its peak (I_peak_) and at the end of the pulse (I_ss_) from experiments such as (E). Averaged I-V relationships of I_A_ from experiments such as (F) show no sustained outward currents (right). (H) Averaged I_Kv4.1_ and I_A_ from LF-GCs are superimposed. (right) I_sus_/I_peak_ for I_Kv4.1_ (red) and I_A_ (grey) of LF-GCs. (Ia) The impact of the Kv4.1 antibody internal on K^+^ current during the bath application of 5 mM 4-AP and 40 µM 4-AP. (Ib) Summary bar graphs showing inhibition of I/I_max_ by Kv4.1 antibody in the presence of 5 mM 4-AP (n = 4) or 40 µM 4-AP (n = 4) in the bathing solution. Error bars indicate SEM. Values indicate mean ± SEM.

We therefore referred to I4-AP in the presence of Kv4.1 antibody as IA. Current-voltage relationships for IKv4.1 and IA are shown in Fig. 6G. To demonstrate the difference between IKv4.1 and IA more clearly, averaged trace for IKv4.1 (obtained from 7 cells) and that for IA (obtained from 10 cells) obtained at +30 mV were superimposed (Fig. 6H). Time-to-peak was not significantly different (at +30 mV, 4.73 ± 0.76 ms for IKv4.1 and 5.31 ± 0.41 ms for IA, *P* > 0.05), suggesting that the activation kinetics for IKv4.1 and IA are similar. In contrast, inactivation kinetics for IKv4.1 was distinctively slower than that for IA. As a result, Iss was larger for IKv4.1 than IA despite that Ipeak was larger for IA than IKv4.1, and the ratio of Iss to Ipeak (Iss/Ipeak) for IKv4.1 was significantly larger than that for IA (0.47 ± 0.11 for IKv4.1, 0.059 ± 0.009 for IA, *P* < 0.01, Fig. 6H). These results suggest that large sustained currents for I4-AP in LF-GCs (Fig. 6C) are attributable to slowly inactivating IKv4.1. Sustained outward currents due to slow inactivation kinetics may support the roles of Kv4.1 as a frequency regulator during repetitive activity.

Since the slow inactivation kinetics of IKv4.1 in GCs resemble D-type K^+^ current (ID) (Storm, 1988; Beck et al., 1997), which shows high sensitivity to 4-AP, we examined 4-AP sensitivity of Kv4.1 currents. To this end, the effect of Kv4.1 antibody on outward K^+^ currents was tested in the presence of a low and a high concentration (40 µM and 5 mM) of 4-AP. Kv4.1 antibody reduced outward currents in the presence of 40 µM, but not 5 mM 4-AP (Figs. 6Ia, b). The fact that a high concentration of 4-AP is required to inhibit IKv4.1 suggests that IKv4.1 is distinctive from ID.

### Kv4.1 expression is developmentally regulated

Rin is inversely correlated with dendritic length which increases with maturation (Dieni et al., 2013). Contribution of Kv4.1 currents to frequency regulation exclusively in LF-GCs where Rin is mostly lower than 200 MΩ may suggest that Kv4.1 expression is developmentally regulated. To investigate this possibility, we examined Kv4.1 expression in the DG of 3-week-old mice and found that Kv4.1 signal was much weaker compared to that in 8-week-old adult mice (Figs. 7A, B). We also noticed a difference in the intensity of Kv4.1 signals between the outer and inner GCL, with cells located at the hilar border showing lower expression (Figs. 7A, B). When newly generated neurons were identified with the doublecortin (DCX) antibody, Kv4.1 signals were hardly found in DCX^+^ cells located in the deep layer both in 8-week-old and 3-week-old mice (Fig. 7B), supporting the maturation-dependent change in Kv4.1 expression. In contrast to Kv4.1, Kv4.2 signal in the DG was not different between 3- and 8-week-old mice (Fig. 7C). The maturation-dependent expression of Kv4.1 was further confirmed by western blot analysis (Fig. 7D).

**Figure 7.**
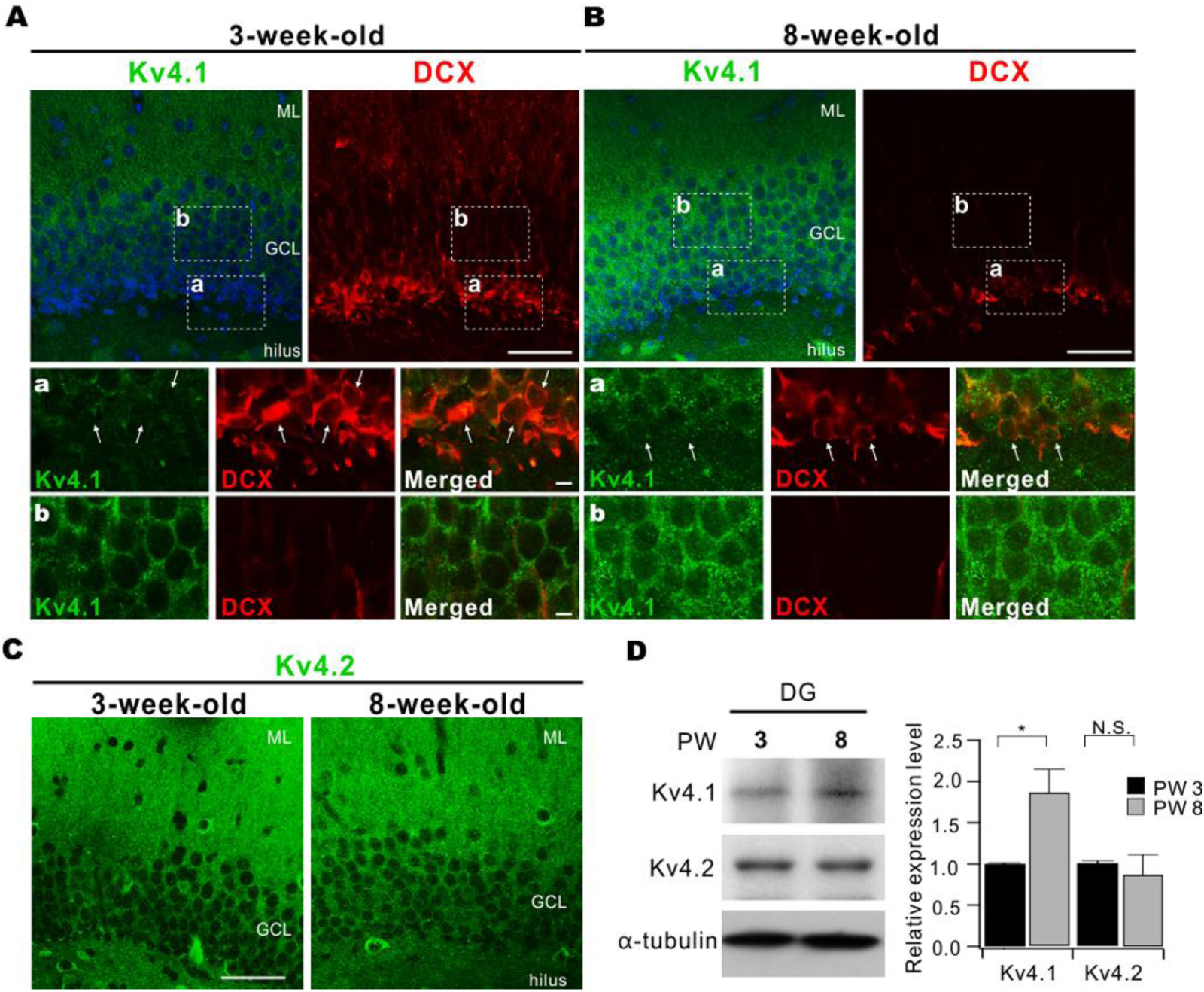
Expression levels of Kv4.1 in DG increase with development. (A, B) Co-immunostaining of Kv4.1 (green) and doublecortin (DCX, red) in 8-week-old (A) and 3-week-old (B) mice. Scale bar, 50 µm. (bottom, a-b) High magnification images of the area corresponding to the dashed box shown in top panels. Scale bar, 5 µm.(C) Immunostaining of DG with Kv4.2 antibody (green) in 8-week-old (left) and 3-week-old (right) mice. Scale bar, 50 µm. (D) Western blot analysis for Kv4.1 and Kv4.2 in DG from 3- and 8-week-old mice. Bar graphs indicate the mean ± SEM (n = 3). GCL, granule cell layer; ML, molecular layer.

### Kv4.1 contributes to sparse firing of synaptically driven APs in LF-GCs

We then examined whether Kv4.1 influences synaptically-evoked membrane events. We applied burst stimulation (5 stimuli at 50 Hz) through a stimulating electrode placed in the middle of the molecular layer to stimulate afferent fibers from the entorhinal cortex, and recorded excitatory postsynaptic currents (EPSCs) or excitatory postsynaptic potentials (EPSPs) in LF-GCs (Fig. 8A). Application of the Kv4.1 antibody did not affect EPSCs (Figs. 8B-D), but gradually increased the amplitude of EPSPs (Figs. 8E, F). Accordingly, the increased EPSPs led to an increase in the number of APs evoked by synaptic stimulation (Fig. 8G). To quantify this effect, we obtained the relationships between stimulus intensity (SI) and AP probability and found that the SI required for the AP probability of 50% (SI50) significantly decreased after blocking Kv4.1 (Fig. 8H). In contrast, the Kv4.1 antibody had no effect on EPSPs in GCs with Rin lower than 200 MΩ depleted of Kv4.1 transcripts using shKv4.1 (Fig. 8I). We also confirmed that the Kv4.1 antibody did not affect the responsiveness to synaptic stimulation in CA1-PCs (Fig. 8J). Taken together, these results suggest that Kv4.1 suppresses EPSPs, and thus lowers the spiking probability of LF-GCs, which may contribute to the sparse activity of the DG.

**Figure 8.**
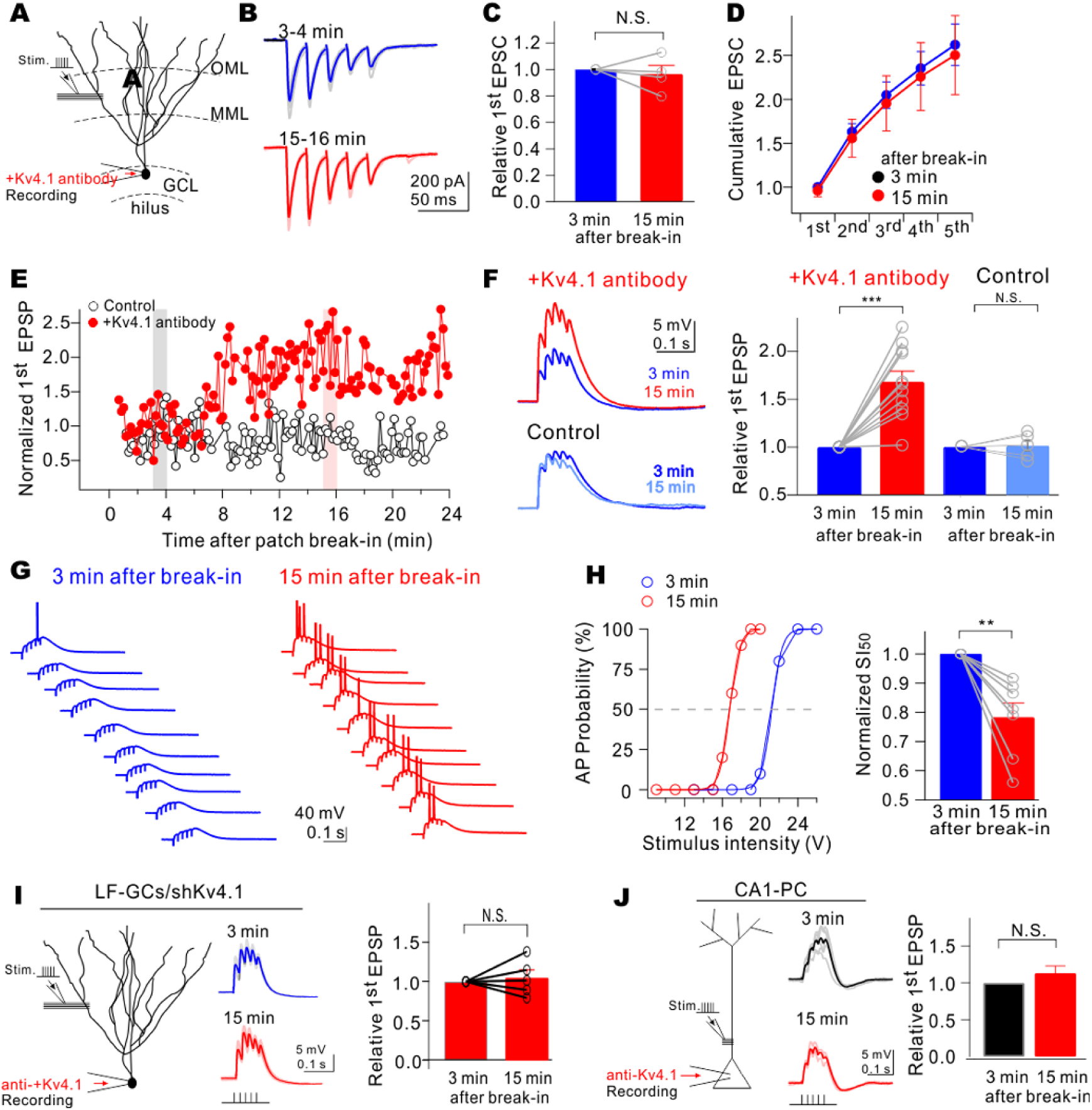
Kv4.1 inhibition increases AP outputs induced by perforant path stimulation in LF-GCs. (A) Schematic diagram of whole-cell recording in LF-GCs with extracellular synaptic stimulation at proximal dendrites. (B) Representative traces of average cumulative EPSCs at 3 min (blue) and 15 min (red) after break-in with the pipette solution containing Kv4.1 antibody. (C-D) Pooled data of normalized 1^st^ EPSC (C) and cumulative EPSC (D) at 3 min (n = 4) and 15 min (n = 4) after break-in with Kv4.1 antibody in the pipette from (B). (E) Time course showing the change of 1^st^ EPSP of mature GCs to 50 Hz stimulation at proximal dendrite with (red) or without (grey) Kv4.1 antibody in the pipette. The amplitude of 1^st^ EPSP is normalized to that at t=0. Insets show average EPSP traces from the time points marked by grey and red shaded boxes. (F) Representative trace of average summated EPSPs after 3 min and 15 min after break-in with Kv4.1 antibody (upper) or vehicle (control, lower) in the pipette. (Right) Bar graph, pooled data of normalized 1^st^ EPSP at 3 min (n = 11) and 15 min (n = 11) after break-in with Kv4.1 antibody in the pipette from (E). Paired t*-*test. \*\*\**P* <0.001. (G) Representative traces from spiking probability measurements. (H, left) AP probability for one GC plotted against stimulus intensity at 3 min (blue) and 15 min (red) after break-in with Kv4.1 antibody in the pipette (sigmoidal fit). (H, right) Summary bar graph illustrating that Kv4.1 antibody treatment decreased the stimulus intensity of 50% spike probability point (SI_50_) (n = 7). Paired t*-*test. \*\**P* < 0.01. (I) Effects of Kv4.1 antibody on EPSPs in GCs depleted with Kv4.1 transcript. Sample traces show EPSPs recorded 3 and 15 min after patch break-in (left). Intracellular perfusion of Kv4.1 antibody had no effect on EPSPs in GCs (R_in_ lower than 200 M) infected with AAV-shKv4.1 (right). (J) Schematic diagram of EPSP recordings in CA1-PCs and sample traces of EPSPs measured in CA1-PCs during 3-4 and 15-16 min after the patch break-in (left). Summary of normalized 1^st^ EPSP at 3 and 15 min (n = 11) after break-in with Kv4.1 antibody in CA1-PCs (right). N. S., *P* > 0.05.

### Impaired pattern separation in mice depleted of Kv4.1 transcripts in DG

Sparse activity is one of the key features that enable the DG to function as a pattern-separator (Treves and Rolls, 1994). To investigate the possibility that Kv4.1 currents in LF-GCs contribute to sparse activity of DG networks and thus to pattern separation, we used a contextual fear conditioning paradigm (McHugh et al., 2007) and tested whether mice with their Kv4.1 transcripts knocked down in GCs exhibit an impairment in discriminating similar contexts. We confirmed that three weeks after the injection of shKv4.1, the expression of GFP was localized in the GCL in the DG and mossy fiber (MF) (Fig. 9A). We first examined the locomotor activity and the anxiety levels of mice using the open field test and elevated plus maze test. Both the shCtrl-injected controls and shKv4.1-injected mice showed similar exploratory patterns as they moved the same distance in the open filed box, and they spent a similar amount of time in the center zone (Fig. 9B). In addition, both groups spent the same amount of time in the open and closed arms of elevated plus maze (Fig. 9C).

**Figure 9.**
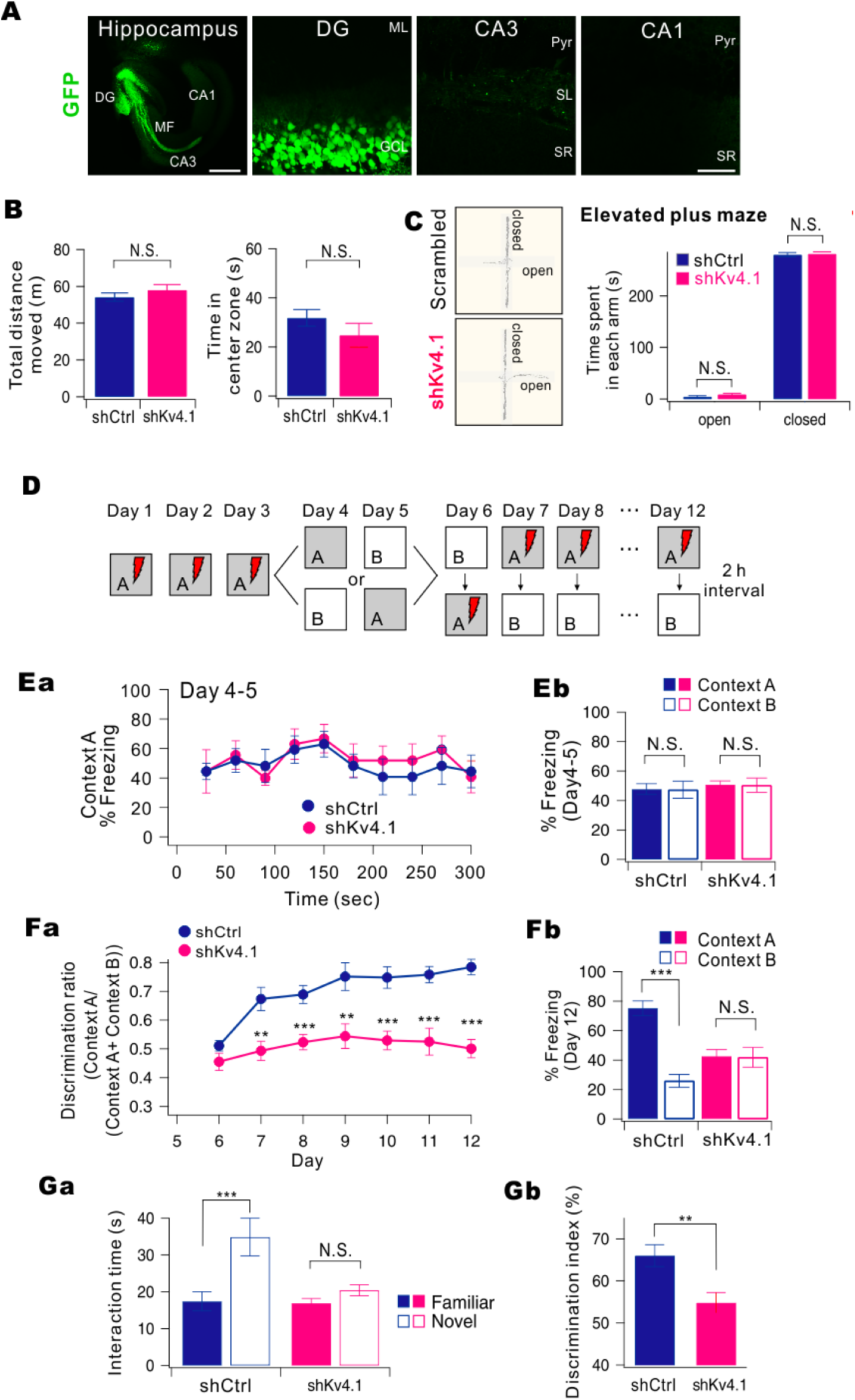
Knock-down of Kv4.1 in DG impairs pattern separation in mice. (A) Sample images of a 12-week-old mouse that had received a stereotaxic injection of AAV-shKv4.1. GFP expression was restricted to the DG area. Left scale bar, 500 µm. Right scale bar, 50 µm. (B-C) DG-specific knock-down of Kv4.1 did not affect the general activity and anxiety levels of mice. (B) Bar graph of total distance traveled for open field box (left) and time spent in the center zone for initial 5 min (right) in shCtrl-injected (black, n = 9) and shKv4.1-injected mice (red, n = 9), *P* > 0.5. (C) Representative tracking trace of the elevated plus maze (left). Right, Bar graph of time spent in each arm for control (dark blue, n = 9) and shKv4.1-injected mice (pink, n = 9). (D) Experimental procedure for pattern separation in 10-week-old mice. (Ea) On day 4 to 5, the kinetics of freezing across the 5 min test in context A. (Eb) Percentage of freezing in context A (filled bars) and context B (open bars) during day 4 to 5. Control (dark blue, n = 9) and shKv4.1 (pink, n = 9) mice displayed equal amounts of freezing inboth contexts A and B. (Fa) Time course of the discrimination ratio in control (dark blue, n = 9) and shKv4.1-injected (pink, n = 9) mice during day 6 to 12. Statistical significance for each day was tested using ANOVA (\**P*< 0.05; \*\**P* < 0.01; *** *P* < 0.001). (Fb) Percentage of freezing in context A (filled bars) and context B (open bars) for control (black, n = 9) and shKv4.1-injected (red, n = 9) mice on day 12. Freezing levels were compared between the two contexts for each group (N.S., no statistical significance; \*\*\**P* < 0.001). (Ga) Time spent interacting with familiar (filled bars) and novel (open bars) objects in shCtrl-(n = 9) and shKv4.1-injected (n =9) mice are summarized. (Gb) Bar graphs show the discrimination index of each group for the novel object.

For behavioral pattern separation analysis, shCtrl-injected controls and shKv4.1-injected mice were tested in a contextual fear conditioning task for the first three days using a context (referred to as ‘context A’) in which animals received a single foot shock 3 min after being introduced to the context (Fig. 9D). On days 4 and 5, we prepared another context similar to the context A and named it ‘context B’. Context B was composed of an identical metal grid floor but had an odor (1% acetic acid), dimmer lighting (50% of context A), and a sloped floor (15° angle). Mice in control and shKv4.1 groups were each divided into two subgroups. One subgroup visited context A on day 4 and context B on day 5, and the other subgroup visited context B on day 4 and context A on day 5. Neither group received a foot shock in either context on days 4 and 5, and freezing was evaluated for 5 min. On days 4 and 5, both the control and shKv4.1-injected groups showed similar freezing levels during the 5-min test in context A (Fig. 9Ea) and appeared to generalize between the two similar contexts (Fig. 9Eb, two-way ANOVA, group, *F*(1,_3_2) = 0.46, *P* = 0.50, context: *F*(1,_3_2) = 0.01, *P* = 0.93, group × context: *F*(1,3_6)_ = 0.00, *P* = 1.00).

The mice were subsequently trained to discriminate between the two contexts by visiting both contexts daily for 7 days with a 2-h interval between the contexts (from day 6 to 12), always receiving a foot shock 180 s after being placed in context A but not in context B. Daily discrimination ratio was calculated by determining the ratio of freezing during the 180-s period in context A to total freezing during the two visits (A and B). On day 6, both groups did not distinguish between the contexts (Fig. 9Fa; group, *F*_(1,32)_= 0.13, *P* = 0.72; context, *F*_(1,32)_= 0.21, *P* = 0.64, two-way ANOVA), and thus the discrimination ratio was approximately 0.5. As the experiment progressed, control mice began to discriminate context B from context A, and the discrimination ratio increased accordingly. However, shKv4.1-injected mice exhibited high levels of freezing in the shock-free context until day 12 (Fig. 9Fb, group: *F*_(1,16)_ = 54.55, *P* < 0.001, group x days: *F*_(6,96)_ = 2.49, *P* < 0.028, repeated measure two-way ANOVA). In Fig. 9Fb, the freezing levels on day 12 are compared between the two contexts for each group (control, *P* < 0.001; shKv4.1, *P* = 0.78; paired t-test), showing deficits in the rapid acquisition of contextual discrimination.

We further examined the behavioral role of Kv4.1 in discrimination of similar objects using the novel object recognition test. During the familiarization phase, both control and shKv4.1-injected mice were allowed to explore two identical sample objects for 10 min. After a 24-h delay, one of the familiar objects was replaced by a similar novel object, and mice were tested for discriminating the novel object from a familiar one. The control group showed a capability of discriminating similar objects and explored the novel object longer (Fig. 9Ga). In contrast, shKv4.1-injected mice spent a similar amount of time exploring familiar and novel objects. Consistent with a lack of preference for the novel object, discrimination index ((time spent exploring the novel object / total object exploration time) x 100%) was significantly reduced in shKv4.1-injected mice (Fig. 9Gb). Collectively, these results suggest that the hyperexcitability of GCs induced by Kv4.1 knockdown specifically impairs pattern separation without affecting the locomotor activity or the general anxiety levels.

## Discussion

In the present study, we identified Kv4.1 as a specific mechanism responsible for low frequency firing of mature GCs. We found that subcellular distribution and biophysical properties of Kv4.1 are distinctive from those of Kv4.2 that mediates classical I_A_ (Figs. 1, 6). We also showed that Kv4.1 inhibition increases the amplitude of EPSPs and AP probability in GCs induced by synaptic inputs from the EC (Fig. 8). Importantly, mice lacking Kv4.1 specifically in the DG were significantly impaired in conditional freezing between similar contexts (Fig. 9), suggesting that low frequency firing of mature GCs is crucial for pattern separation.

### Changes in intrinsic excitability and firing behavior during maturation of GCs

The DG is where adult neurogenesis occurs. Changes in excitation/inhibition balance and intrinsic excitability are key components that occur during maturation of GCs. In contrast to the rigorous studies on maturation-dependent synaptic changes (Marin-Burgin et al., 2012; Dieni et al., 2013), maturation-dependent changes in intrinsic excitability, in particular the mechanisms underlying firing frequency changes, are not fully understood. It is generally believed that hyperpolarization of RMP occurs in parallel with the decrease in R_in_ during maturation, resulting in low frequency firing in mature GCs. Inwardly rectifying potassium channels (K_ir_) (Mongiat et al., 2009) and GABA_B_ receptor-mediated GIRK channels (Gonzalez et al., 2018) were suggested to underlie maturation-dependent decrease in firing frequency in adult-born GCs. GIRK current appears in developing GCs with R_in_ between 1 and 2 GΩ which corresponds to ∼3 weeks of age (Gonzalez et al., 2018), while K_ir_ expression increases between 4 to 7 weeks (Mongiat et al., 2009). In the present study, we show that RMP hyperpolarization occurs steeply as R_in_ decreases lower than 300 MΩ (Fig. 3B), while abrupt decrease in firing frequency due to Kv4.1 occurs when R_in_ decreases lower than 200 MΩ (Fig. 4G), suggesting that Kv4.1 expression occurs later than GIRK or K_ir_ expression, at the very last stage of maturation.

Kv4.1 may play distinctive roles from K_ir_ or GIRK channels in lowering the excitability of GCs. We showed that Kv4.1 inhibition increases the gain of F-I curve (Fig. 4E), whereas inhibition of K_ir_ using BaCl_2_ or GIRK using a GTP-free internal solution shifted the F-I curve to the left with little change in its gain (Mongiat et al., 2009). Considering that K_ir_ and GIRK are active at RMP, this difference may suggest that inhibition of background leak currents is capable of reducing threshold currents, but does not affect the gain of input-output relationship significantly. In contrast, Kv4.1 that is activated at depolarized potentials and remains active during repetitive APs (Fig. 6) can regulate the I-O gain.

Brunner et al. (2014) monitored various intrinsic excitability parameters in adult-born GCs from 3 to 10 weeks of age and found that there are two functionally distinct groups in adult-born GCs with respect to the I-O gain. Youngest GCs (3–5 weeks old) are very sensitive to input (S-group, high I-O gain) and oldest GCs (10 weeks old) are less sensitive and respond linearly to input (L-group, low input-output gain). Interestingly, the transition from S-group to L-group occurs randomly between 5 and 9 weeks without correlation with the age of the cells or R_in_ (Brunner et al., 2014). These observations were consistent with the idea that the mechanism responsible for decreasing the input-output gain is different from the mechanism that underlies maturation-dependent decrease in R_in_ or RMP, but they could not identify what the mechanism was. We provide a line of evidence that Kv4.1 is responsible for the low I-O gain in mature GCs. To our knowledge, our study is the first to identify maturation-dependent changes in voltage-gated K_^+^_ channels that can specifically control firing probability.

### Distinctive properties of Kv4.1 compared to other Kv4 currents

Kv4.1 has similar electrophysiological properties to those of Kv4.2 and Kv4.3 when expressed in heterologous expression systems (Pak et al., 1991), showing rapid inactivation kinetics. In a recent study where a role of Kv4.1 in the suprachiasmatic nucleus was suggested, Kv4.1 currents were characterized as A-type K_^+^_ currents (Hermanstyne et al., 2017). We found that biophysical properties of Kv4.1 characterized using Kv4.1 antibody in GCs showed a slowly inactivating component (Fig. 6E), which are distinctive from those of classical A-type K^+^ currents. The reason for the observed discrepancy between the biophysical characteristics of I_Kv4.1_ in expression systems and GCs is currently unclear. It is plausible that Kv4.1 activity in native neurons might be regulated through diverse modulatory mechanisms including phosphorylation and accessory proteins, which may be absent in *in vitro* expression systems.

We found that there was an obvious contrast between the distribution of Kv4.1 and Kv4.2 (Figs. 1B, C) in the hippocampus, but the mechanisms and the physiological significance of the distinctive distribution remain to be investigated. In CA1-PCs, I_A_ was shown to be mediated by Kv4.2, mainly regulating AP repolarization phase (Kim et al., 2005; Chen et al., 2006; Kim et al., 2007). The role of I_A_ in the regulation of dendritic excitability in GCs was proposed from the experiment using 4-AP puff around 50 μm from the soma (Lopez-Rojas et al., 2016), but the molecular identity of ion channels underlying 4-AP effect remains to be uncovered. We showed that Kv4.1 inhibition increases EPSP amplitudes (Fig. 8). Considering that 4-AP can inhibit Kv4.1 as well as Kv4.2, interpretation of 4-AP effects needs to be revisited. Contribution of Kv4.3 to 4-AP effects is also likely since the Kv4.3 subunit is highly expressed in the molecular layer of the DG same as the Kv4.2 subunit (Clark et al., 2008). Future studies are required to determine the relative roles of Kv4 family channels in dendritic excitability of GCs.

### Physiological and pathophysiological implications of Kv4.1 in mature GCs

It has been postulated that pattern separation reflects non-overlapping representations of memories on the network (O’Reilly and McClelland, 1994), and thus sparse network activity is prerequisite for pattern separation. Frequency regulation of GCs by Kv4.1 may play a key role in limiting the network activity not only in the DG but also in the CA3 region, since a single MF can induce firing of postsynaptic CA3-PCs only if the inputs are at high frequency (Henze et al., 2002; Bischofberger et al., 2006), whereas low frequency MF inputs may rather have inhibitory influences to CA3 network via feed-forward interneurons (Mori et al., 2004). We also noted that that Kv4.1 is highly expressed also in CA3 (Fig. 1A), where the network activity is also sparse *in vivo* (Leutgeb et al., 2004; GoodSmith et al., 2017; Senzai and Buzsaki, 2017). The role of Kv4.1 in CA3-PCs needs to be investigated.

In the present study, we did not exclude the possibility that interneurons in the DG might also be infected with AAV-shKv4.1, but we confirmed that Kv4.1 antibody has no effect on spike frequency of interneurons in the DG (data not shown). In accordance with our observation, Kv4.1 transcripts were not detected in hippocampal interneurons (Serodio and Rudy, 1998). Thus, our results suggest that impaired pattern separation caused by the knockdown of Kv4.1 mainly stem from abnormally enhanced excitability in mature GCs.

Mice with ablated adult neurogenesis showed impaired pattern separation when stimuli were presented with little spatial separation (Clelland et al., 2009), while enhanced adult hippocampal neurogenesis improved pattern separation (Sahay et al., 2011). In addition, a previous study has shown that transgenic mice where output of mature GCs was specifically inhibited exhibited enhanced or normal pattern separation function (Nakashiba et al., 2012). It was thus suggested that pattern separation seems to be mediated mainly by young GCs rather than mature GCs. We, however, showed that hyperexcitability of mature GCs caused by knockdown of Kv4.1 induces impaired pattern separation, implying that the low excitability of mature GCs is critical for pattern separation. Our results do not necessarily mean that mature GCs play a key role in pattern separation. Possibly, hyperexcitability induces greater participation of mature GCs in DG network and hampers young GCs from discriminating similar experiences. A possibility that mature GCs are not passive bystanders but principal performer in DG functions was also proposed (Lopez-Rojas and Kreutz, 2016). Several recent papers indirectly suggest that mature GCs participate in learning and memory formation. Active GCs identified during spatial exploration corresponded to morphologically mature GCs (Diamantaki et al., 2016), and the engram-dentate neurons have electrophysiological characteristics of mature GCs (R_in_ of 100 MΩ) (Ryan et al., 2015). DG network composed of mature GCs and more excitable and plastic young GCs may work together to accomplish pattern separation.

Although the roles of Kv4.1 in brain functions have not been well investigated, possible involvement of Kv4.1 in neuropsychiatric diseases has been noted. Kv4.1 is one of the genes exhibiting a significant number of variants in schizophrenia patients (Piton et al., 2011), and also suggested as a candidate gene for a new neurological syndrome with mental retardation, choreoathetosis, and abnormal behavior which has been mapped to chromosome Xp11 (Reyniers et al., 1999). Combined with our experimental data, Kv4.1 can be a potential target for the intervention of cognitive deficit or other neuropsychiatric diseases.

## Acknowledgements

This research was supported by the National Research Foundation of Korea (NRF) grants (2010-0027941 and 2017R1A2B2010186 to W.-K. Ho, and 2017R1A6A3A11032599 to K.-R. Kim) funded by the Korean Ministry of Science and ICT.

## Author contributions

S.H.L. and W.-K.H. developed the study concept and design. K.R.K., S.Y.L., Y. K., M.J.K., S.L., and H.J.J performed experiments and collected data under the supervision of Y.H.S., J.S.K., and W.-K.H. S.H.L., M.-H.K., H. C., and W.-K.H. analyzed and interpreted the data, prepared the figures and wrote the manuscript. All authors contributed to and approved the manuscript.

